# Identification of novel senolytic compounds from natural food sources

**DOI:** 10.1101/2022.05.12.491721

**Authors:** Tesko Chaganti, Brahmaiah Pendyala

**Affiliations:** Vincent Massey Secondary School, Windsor, ON, Canada; Department of Agricultural and Environmental Sciences, Food Science Program, College of Agriculture, Tennessee State University, Nashville, TN, United States

**Author notes:** Corresponding author **Brahmaiah Pendyala, Ph.D.**, Research Scientist.

**Keywords:** PI3Kγ, senolytic compounds, natural compounds, molecular docking, age associated diseases, molecular dynamics

## Abstract

More than 23% of today’s population suffers from age-associated diseases such as arthritis, cancer, heart disease, and more. The ongoing economic impact of these diseases has been in the billions of dollars worldwide with no clear solution to date. This study addresses the underlying cause of these diseases by identifying the compounds that potentially eliminate senescent cells. Existing senolytic drugs are not abundantly found in nature, reducing accessibility. Hence, over 70,000 natural compounds available in the Canadian Food Database were used to screen penitential senolytic compounds that block PI3Kγ, reactivating apoptotic processes in senescent cells. Molecular docking results revealed 23 natural compounds that blocks the PI3Kγ. Out of 23 compounds, Cianidanol, Ellagic acid, Eriodictyol, Kaempferol and Cyanidin were found abundantly in food sources range from 85 to 735 mg/100 g. These compounds are up to 46 times more abundant in foods than proven senolytic drug Fisetin. Further, molecular dynamics results showed ligand stability for 4 nanoseconds with PI3Kγ. The five compounds are proven to eliminate cancerous cells, have the potential to prevent age-related diseases, and could even slow down natural aging.

## Introduction

Canada spends nearly half of its health care expenditure on people in retirement (Globerman, 2021). Many of these people suffer from age-associated diseases such as arthritis, cardiovascular disease, cancer, osteoporosis, as well as numerous other diseases (von Kobbe, 2019). Most of these diseases have no cure and have few preventatives. A harmful cell type known as senescent cells are considered pivotal to not only the aging process, but also the inception of age-associated diseases (Tuttle et al., 2019). Originally these cells were healthy, but due to certain circumstances, such as high stress or shortened telomeres, the cells undergo an irreversible process of cell cycle arrest and can become senescent (Childs et al,. 2015). This means that these cells stop dividing, and stop being useful to the body. Senescent cells irritate nearby cells by releasing Senescence Associated Secretory Phenotypes, stimulating them into becoming senescent, or even worse, cancerous (von Kobbe, 2019). By removing senescent cells from the body, people can delay the speed at which they age and reduce the likelihood of acquiring chronic diseases (Tuttle et al., 2019). Fisetin, a proven senolytic drug (Yousefzadeh et al., 2018), blocks an anti-apoptotic pathway, phosphatidylinositol 3-kinase/protein kinase B (PI3K-AKT) (Chien et al., 2010). The signaling pathway is responsible for inhibiting apoptosis related mechanisms that would otherwise cause the senescent cell to undergo apoptosis (Gasek et al., 2021). Specifically, Fisetin inhibits the activator of this pathway, PI3K, thus causing apoptosis in senescent cells (Lim et al., 2015). While Fisetin expresses senolytic activity, it is not abundantly found in food (0.16 mg/g in strawberries (Kahn et al., 2013)). Quercetin is abundant in foods (149 mg/g in evening primrose (Canadian Food Database: [FooDB], 2020)), but its senolytic activity is debated (Hwang et al., 2018). The aim of this project is to discover novel senolytic compounds that are highly abundant in everyday foods such as fruits and vegetables which perform the same tasks as the proven senolytic drug Fisetin.

## Materials and Methods

### Data sources and primary screening

FooDB was used, which is the largest food database globally with 70,962 natural compounds (FooDB, n.d), to discover compounds that have the potential to function as senolytic drugs. To generate a shortlist of compounds from the main database, compounds (ligands) that were detected and quantified in foods were filtered. Then, all hydrocarbons, derivatives, and ligands with a molecular weight of greater than 600g/mol were removed. The remaining ligand files and the PI3Kγ receptor file (PDB ID: 1E8Y) were downloaded from PubChem and the Protein Data Bank (PDB) databases, respectively, and Visual Studio Code with Python was used to export the names and atomic masses of the ligands using their PubChem IDs to Google Sheets from PubChem API. Compound food sources were procured from FooDB.

### Molecular docking

The programs AutoDock Tools and Open Babel were used to prepare the necessary files for docking. AutoDock Vina was used on a Virtual Machine running Ubuntu via Oracle VirtualBox, and the downloaded ligands were docked to the PI3Kγ receptor, generating binding energy values. Next, compounds with a binding energy of greater than -8kCal/mol were excluded, and the ratio of atomic mass to the binding energy was calculated. The compounds with the lowest mass ratio were then selected to manually screen using PyMol, an open-source python-based molecule visualization program, to identify the compounds binding to key amino acids (E883, D841, E880, V882, D964 (Lim et al., 2015)) in the ATP binding site of PI3K, inhibiting it. The sources of compounds successfully blocking the PI3Kγ receptor were obtained using FooDB.

### Molecular dynamics

Molecular Dynamics was used to examine protein-ligand interactions in a simulated environment of the human body. The files necessary for molecular dynamics were collected and prepared using Visual Molecular Dynamics (VMD) (Humphrey et al., 1996) and CHARMM-GUI (Jo et al., 1995). Molecular dynamics simulations were performed on the docked protein-ligand complexes using NAMD (Phillips et al., 2005). Finally, VMD was used to analyze the outputs of the simulation and generate Root-Mean-Squared Deviation (RMSD) values.

## Results and Discussion

### Primary screening

The search criteria was to find highly abundant food compounds that can inhibit PI3Kγ. So, all unquantified and undetected compounds were filtered from FooDB. Unquantified refers to compounds that were detected in food, with extremely low concentrations that are not possible to measure and undetected compounds were not found to be in foods. 3571 compounds remained out of 70,964.

Next, hydrocarbon compounds, derivative compounds, and compounds with a molecular weight of more than 600 g/mol were removed, resulting in 414 compounds. This is because hydrocarbons consist of long chains of atoms that are unreactive and smaller molecules are more efficient in reaching target cells (Vargason et al., 2021).

### Molecular docking

Compounds with lower binding energies have a higher tendency to form strong bonds with the receptor. So, compounds with binding energies less than -8 kCal/mol were kept, and 165 of 414 compounds remained.

The top 75 of 165 compounds with the least mass ratio were selected and visualized to determine if they inhibited PI3Kγ, which resulted in 23 compounds that were found to successfully block the PI3Kγ receptor by binding with it in the ATP binding site.

Finally, all compounds that were found in lower concentrations in food than Fisetin were removed, resulting in five novel compounds (Kaempferol, Eriodictyol, Cyanidin, Cianidanol, Ellagic Acid). The bonds of these compounds with PI3Kγ were shown in Figure 1.

**Figure 1:**
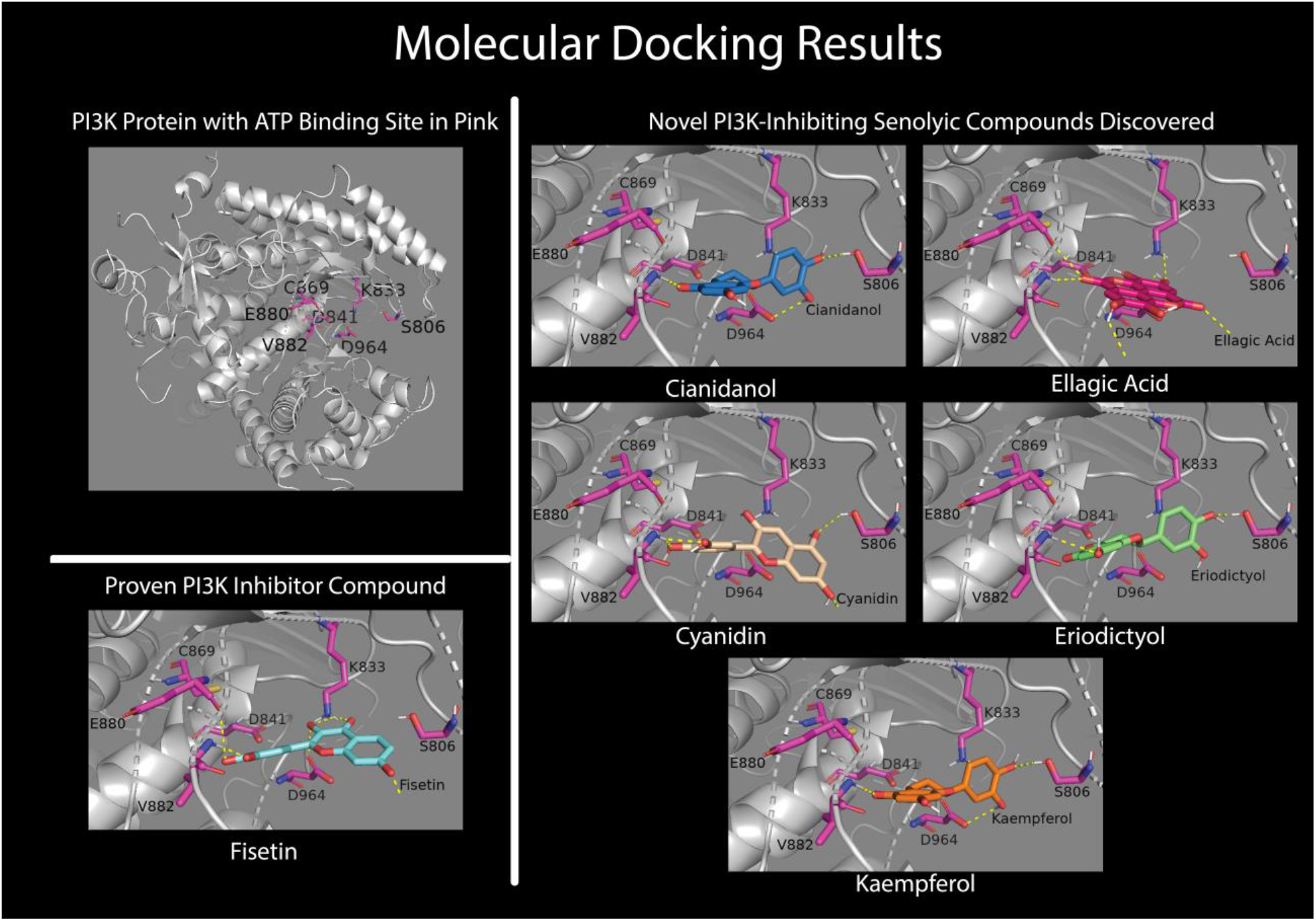
Molecular docking of the five novel PI3K inhibiting compounds compared to Fisetin

### Molecular dynamics

Molecular dynamics simulations conducted on all five compounds showed stability for four nanoseconds with RMSD values less than 2.0 Å (Figure 2). This further establishes that the compounds have strong bonds in active site of PI3K. Out of five compounds tested Ellagic acid and Cianidanol showed better stability than Fisetin (Figure 2). The results of the molecular docking and dynamics simulations indicate that these compounds have a high potential for expressing senolytic activity.

**Figure 2:**
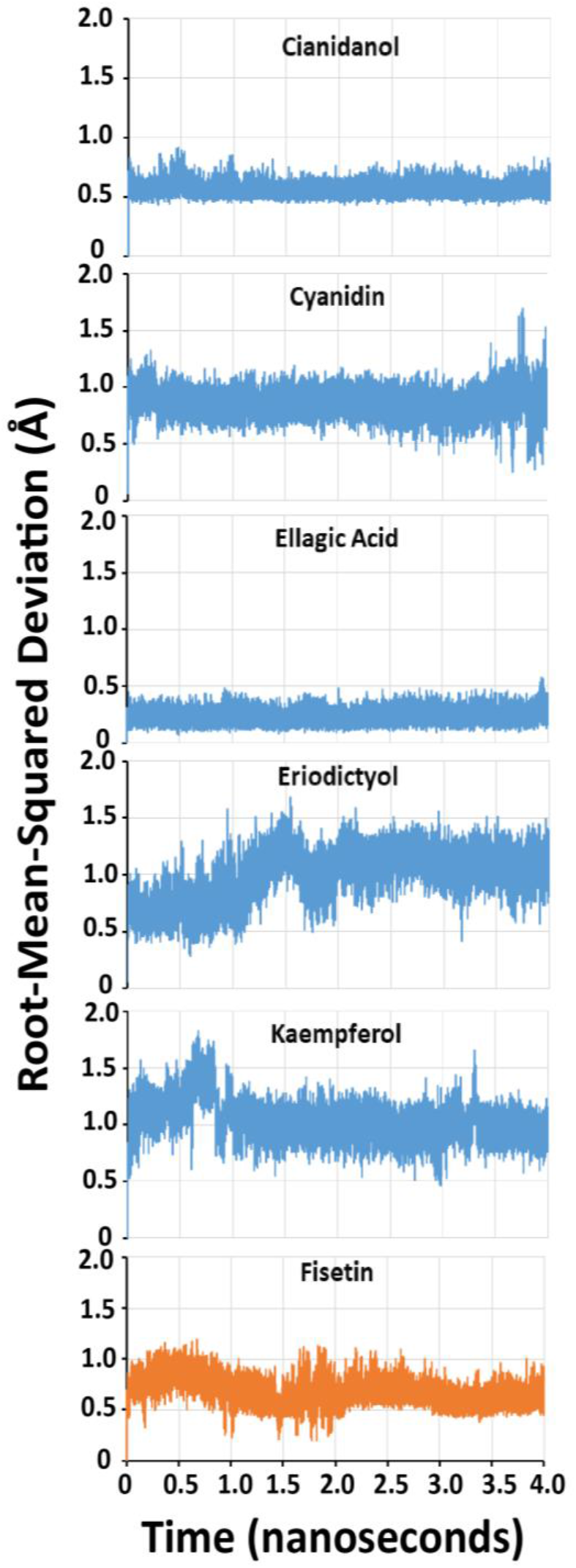
Molecular Dynamics simulations of ligands with PI3K

The identified compounds are found abundantly in herbs and nuts such as Cumin, Oregano, Mexican Oregano, Peppermint, Saffron, Walnuts, and Chestnuts, along with fruits and vegetables like Lemons, Sweet Oranges, Celery, Blackcurrant, Blackberries, and Blueberries (Figure 3).

**Figure 3:**
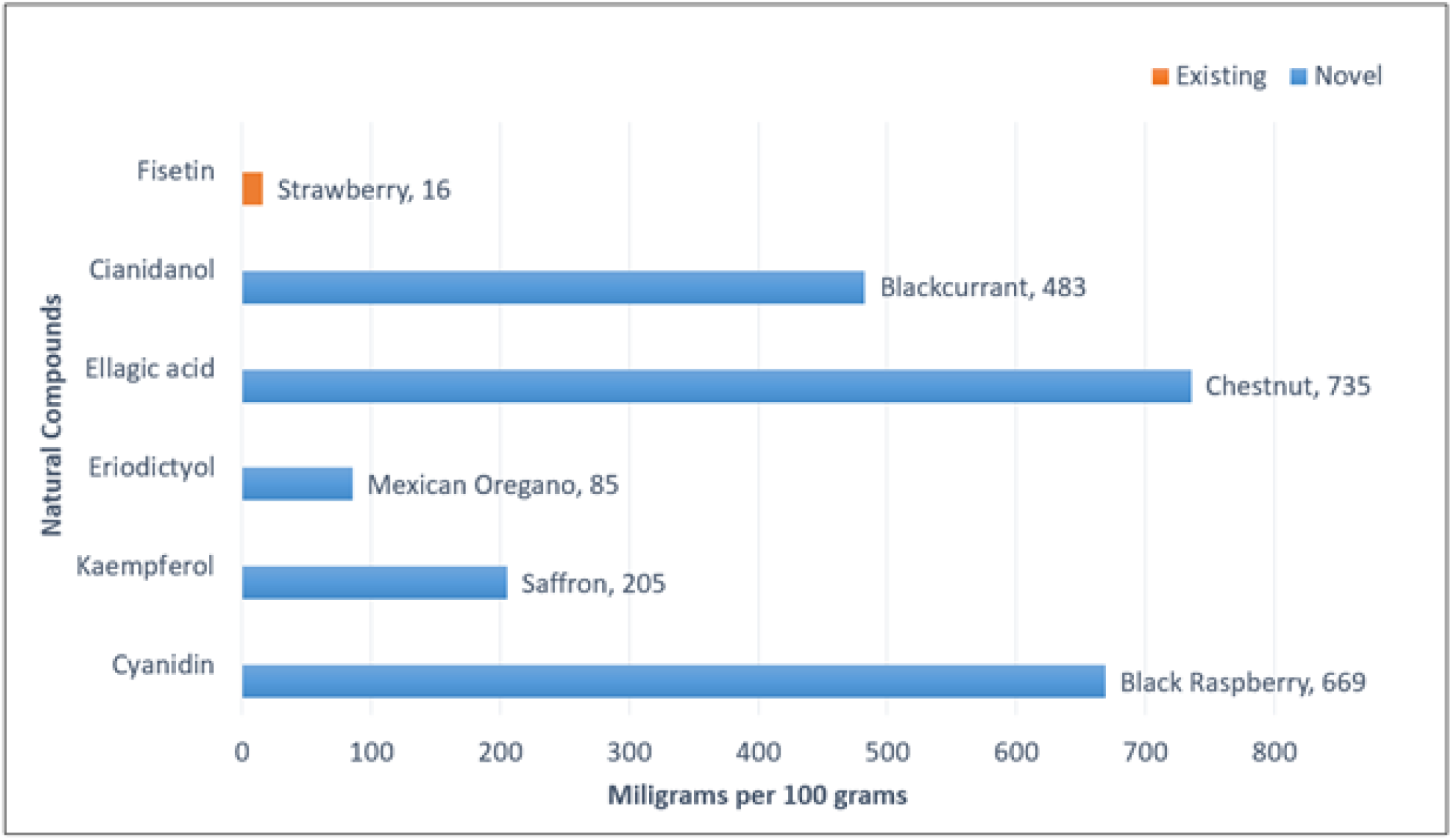
Comparison of novel potential senolytic compounds with Fisetin in major natural food sources.

This project discovered the exact mechanism of five potential novel senolytic compounds that are available in natural food sources. Further, the compounds identified as potential senolytic drugs have known anti-cancer and anti-proliferation properties (Ponte et al., 2021; Wang et al., 2019; Li et al., 2020). In the conformational literature search, Cyanidin, one of the five compounds that was discovered to inhibit PI3K, was already identified as a senolytic compound (Choi et al., 2010). However, there was no mention of the specific protein target PI3K, meaning that in that case, an additional mechanism through which Cyanidin eliminates senescent cells was potentially discovered. Discovery of these compounds as potential senolytics and their specific mechanism to target PI3Kγ is novel and ground-breaking.

## Conclusion

Since all the compounds identified with the potential to selectively eliminate senescent cells from humans are found in edible sources, it is possible that by consuming more or less of specific food items, the aging process could be slowed down. However, further simulations and human cell trials are required to determine ideal dosages for statistically significant effects. These natural compounds have high potential for application in pharmaceuticals and nutraceuticals and could enhance the health of billions of people worldwide.

## Author contributions

TC developed the research idea, analysed the data, performed molecular docking & dynamics, generated figures and wrote the manuscript. BP involved in conceptualization, training TC for molecular docking and dynamics, supervision, reviewing, and editing.

## Notes

### Competing Interest Statement

The authors have declared no competing interest.

## References

Canadian Food Database. (2020, September 17th). Retrieved March 15, 2022, from https://foodb.ca/compounds/FDB011904

Chien, C. S., Shen, K. H., Huang, J. S., Ko, S. C., & Shih, Y. W. (2010). Antimetastatic potential of fisetin involves inactivation of the PI3K/Akt and JNK signaling pathways with downregulation of MMP-2/9 expressions in prostate cancer PC-3 cells. Molecular and cellular biochemistry, 333(1-2), 169–180. https://doi.org/10.1007/s11010-009-0217-z

Childs, B. G., Durik, M., Baker, D. J., & van Deursen, J. M. (2015). Cellular senescence in aging and age-related disease: from mechanisms to therapy. Nature medicine, 21(12), 1424–1435. https://doi.org/10.1038/nm.4000

Choi, M. J., Kim, B. K., Park, K. Y., Yokozawa, T., Song, Y. O., & Cho, E. J. (2010). Anti-aging effects of cyanidin under a stress-induced premature senescence cellular system. Biological & pharmaceutical bulletin, 33(3), 421–426. https://doi.org/10.1248/bpb.33.421

FoodDB. (n.d.). https://foodb.ca

Gasek, N. S., Kuchel, G. A., Kirkland, J. L., & Xu, M. (2021). Strategies for Targeting Senescent Cells in Human Disease. Nature aging, 1(10), 870–879. https://doi.org/10.1038/s43587-021-00121-8

Globerman, S. (2021). Health-care spending may spike by 88 percent due to Canada’s aging population. Fraser institute. https://www.fraserinstitute.org/article/health-care-spending-may-spike-by-88-per-cent-due-to-canadas-aging-population

Humphrey, W., Dalke, A., and Schulten, K. (1996). VMD: visual molecular dynamics. J. Mol. Graph. 14, 33–38. https://doi.org/10.1016/0263-7855(96)00018-5

Hwang, H. V., Tran, D. T., Rebuffatti, M. N., Li, C. S., & Knowlton, A. A. (2018). Investigation of quercetin and hyperoside as senolytics in adult human endothelial cells. PloS one, 13(1), e0190374. https://doi.org/10.1371/journal.pone.0190374

Jo, S., Kim, T., Iyer, V. G., & Im, W. (2008). CHARMM-GUI: a web-based graphical user interface for CHARMM. Journal of computational chemistry, 29(11), 1859–1865. https://doi.org/10.1002/jcc.20945

Khan, N., Syed, D. N., Ahmad, N., & Mukhtar, H. (2013). Fisetin: a dietary antioxidant for health promotion. Antioxidants & redox signaling, 19(2), 151–162. https://doi.org/10.1089/ars.2012.4901

Li, W., Du, Q., Li, X., Zheng, X., Lv, F., Xi, X., Huang, G., Yang, J., & Liu, S. (2020). Eriodictyol Inhibits Proliferation, Metastasis and Induces Apoptosis of Glioma Cells via PI3K/Akt/NF-κB Signaling Pathway. Frontiers in pharmacology, 11, 114. https://doi.org/10.3389/fphar.2020.00114

Lim, J. Y., Lee, J. Y., Byun, B. J., & Kim, S. H. (2015). Fisetin targets phosphatidylinositol-3-kinase and induces apoptosis of human B lymphoma Raji cells. Toxicology reports, 2, 984– 989. https://doi.org/10.1016/j.toxrep.2015.07.004

Phillips, J.C., Braun, R., Wang, W., Gumbart, J., Tajkhorshid, E., Villa, E., Chipot, C., Skeel, R.D., Kalé, L. and Schulten, K. (2005), Scalable molecular dynamics with NAMD. J. Comput. Chem., 26, 1781–1802. https://doi.org/10.1002/jcc.20289

Ponte, L. G. S., Pavan, I. C. B., Mancini, M. C. S., da Silva, L. G. S., Morelli, A. P., Severino, M. B., Bezerra, R. M. N., & Simabuco, F. M. (2021). The Hallmarks of Flavonoids in Cancer. Molecules, 26(7), 2029. https://doi.org/10.3390/molecules26072029

Tuttle, C., Waaijer, M., Slee-Valentijn, M. S., Stijnen, T., Westendorp, R., & Maier, A. B. (2020). Cellular senescence and chronological age in various human tissues: A systematic review and meta-analysis. Aging cell, 19(2), e13083. https://doi.org/10.1111/acel.13083

Vargason, A. M., Anselmo, A. C., & Mitragotri, S. (2021). The evolution of commercial drug delivery technologies. Nature biomedical engineering, 5(9), 951–967. https://doi.org/10.1038/s41551-021-00698-w

von Kobbe C. (2019). Targeting senescent cells: approaches, opportunities, challenges. Aging, 11(24), 12844–12861. https://doi.org/10.18632/aging.102557

Wang, Y., Ren, F., Li, B., Song, Z., Chen, P., & Ouyang, L. (2019). Ellagic acid exerts antitumor effects via the PI3K signaling pathway in endometrial cancer. Journal of Cancer, 10(15), 3303–3314. https://doi.org/10.7150/jca.29738

Yousefzadeh, M. J., Zhu, Y., McGowan, S. J., Angelini, L., Fuhrmann-Stroissnigg, H., Xu, M., Ling, Y. Y., Melos, K. I., Pirtskhalava, T., Inman, C. L., McGuckian, C., Wade, E. A., Kato, J. I., Grassi, D., Wentworth, M., Burd, C. E., Arriaga, E. A., Ladiges, W. L., Tchkonia, T., Kirkland, J. L., … Niedernhofer, L. J. (2018). Fisetin is a senotherapeutic that extends health and lifespan. EBioMedicine, 36, 18–28. https://doi.org/10.1016/j.ebiom.2018.09.015

